# Genomic Characterization of a Dog-Mediated Rabies Outbreak in El Pedregal, Arequipa, Peru

**DOI:** 10.1101/2024.08.21.608982

**Authors:** Renzo Salazar, Kirstyn Brunker, Elvis W. Díaz, Edith Zegarra, Ynes Monroy, Gorky N. Baldarrago, Katty Borrini-Mayorí, Micaela De la Puente-León, Sandeep Kasaragod, Michael Z. Levy, Katie Hampson, Ricardo Castillo-Neyra

## Abstract

**Background:** Rabies, a re-emerging zoonosis with the highest known human case fatality rate, has been largely absent from Peru, except for endemic circulation in the Puno region on the Bolivian border and re-emergence in Arequipa City in 2015, where it has persisted. In 2021, an outbreak occurred in the rapidly expanding city of El Pedregal near Arequipa, followed by more cases in 2022 after nearly a year of epidemiological silence. While currently under control, questions persist regarding the origin of the El Pedregal outbreak and implications for maintaining rabies control in Peru.

**Methods:** We sequenced 25 dog rabies virus (RABV) genomes from the El Pedregal outbreak (n=11) and Arequipa City (n=14) from 2021-2023 using Nanopore sequencing in Peru. Historical genomes from Puno (n=4, 2010-2012) and Arequipa (n=5, 2015-2019), were sequenced using an Illumina approach in the UK. In total, 34 RABV genomes were analyzed, including archived and newly obtained samples. The genomes were analyzed phylogenetically to understand the outbreak’s context and origins.

**Results:** Phylogenomic analysis identified two genetic clusters in El Pedregal: 2021 cases stemmed from a single introduction unrelated to Arequipa cases, while the 2022 sequence suggested a new introduction from Arequipa rather than persistence. In relation to canine RABV diversity in Latin America, all new sequences belonged to a new minor clade, Cosmopolitan Am5, sharing relatives from Bolivia, Argentina, and Brazil.

**Conclusion:** Genomic insights into the El Pedregal outbreak revealed multiple introductions over a 2-year window. Eco-epidemiological conditions, including migratory worker patterns, suggest human-mediated movement drove introductions. Despite outbreak containment, El Pedregal remains at risk of dog-mediated rabies due to ongoing circulation in Arequipa, Puno, and Bolivia. Human-mediated movement of dogs presents a major risk for rabies re-emergence in Peru, jeopardizing regional dog-mediated rabies control. Additional sequence data is needed for comprehensive phylogenetic analyses.

## Introduction

Rabies, a globally prevalent zoonotic disease, has one of the highest fatality rates among both humans and animals, with infections primarily resulting from rabid dog bites (1). The economic burden attributed to rabies surpasses 8 billion USD, with premature death representing a major component (2). In a concerted effort to eliminate human deaths caused by dog-mediated rabies by 2030, a worldwide strategic plan—‘Zero by 30’—was established in 2015 (3). To achieve this global goal, it is crucial to have efficient and well-coordinated local surveillance in place (4). Genomic surveillance, a key tool in molecular epidemiology, provides unique insights into virus dynamics (5,6), spread (7), and control advances (8–10). In the context of dog-mediated rabies, genomic surveillance can complement epidemiological efforts, offering valuable information to comprehend and redirect control strategies during outbreaks (11).

In Peru, dog-mediated human rabies has been mostly controlled (12,13), but there has been continuous rabies virus (RABV) transmission in the dog population of Arequipa City since its detection in 2015 (14,15). In 2021, rabid dogs were detected in El Pedregal, a rapidly growing city in the neighboring province of Caylloma, 2 hours from Arequipa City (16). El Pedregal was built around an irrigation project in the desert; 40 years ago the area was uninhabited, but it has experienced explosive growth and development (17). Given its strategic location and economic opportunities, El Pedregal has become a hub for agricultural employment, with considerable in-migration and commuting from neighboring cities and towns (17,18). The dog rabies outbreak in El Pedregal prompted the local government to implement focused control strategies and strengthen mass dog vaccination campaigns (19). The outbreak has been controlled, but many epidemiological questions remain unanswered. For instance, the outbreak source and the frequency of introductions are yet unknown, the diversity and distribution of circulating virus lineages have not been characterized, and it is unclear to what extent local transmission may be undetected, all questions that can be answered by genomic surveillance (11,20).

Identifying the source of emerging rabies outbreaks is vital for understanding where dog vaccination needs to be strengthened (21) and understanding the potential for dog-mediated RABV re-emergence. While many countries worldwide contend with endemic dog-mediated rabies virus, the case of El Pedregal presents an instance of re-emergence in a previously rabies-free area (and almost dog-mediated rabies-free continent). This case offers valuable insights into the effectiveness of RABV genomic surveillance and its pivotal role in the latter stages of control efforts. It serves as a warning for potential challenges that may arise in other regions globally, as well as specifically informing the current epidemiological situation in Peru and Latin America, highlighting the evolving epidemiological dynamics of the rabies virus as we strive towards and aim to sustain elimination. Therefore, this study aims to elucidate the origin of the outbreak in El Pedregal using epidemiological and genomic data and to characterize the spread of RABV within this unusual epidemiological context.

## Methods

### Study area

This study was conducted in El Pedregal (Fig 1), Majes District, Caylloma Province, Arequipa Department, Peru. Located in the subtropical Coastal desert, approximately 1440 meters above sea level and southwest of Arequipa city (22,23), El Pedregal is home to an estimated population of 70,780 inhabitants (24). The population is mainly composed of migrant populations (17); up to 90% come from nearby cities and regions such as Arequipa, Puno, and Cusco, as well as neighboring towns (17).

**Fig 1.**
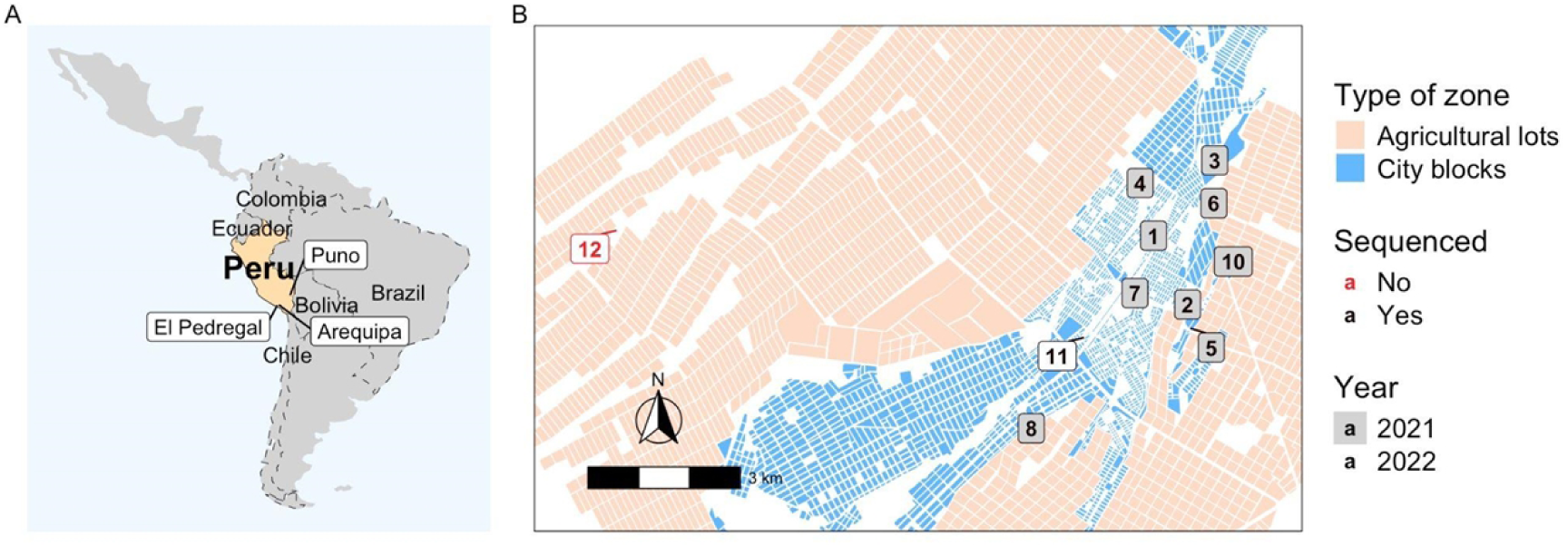
Geographic context of the study area in Peru and zoomed-in map of El Pedregal showing the location of rabies cases during the 2021-2022 outbreak. A) Map of Latin America highlighting Peru, with annotations of locations relevant to the outbreak study. B) Detailed map of El Pedregal, showing the locations of rabies cases in rural and urban areas for 2021 and 2022. Cases are numbered according to the epidemiological timeline.

### The outbreak

On February 9, 2021, the first case of dog rabies was reported in El Pedregal (Fig 1, Table 1). The rabid dog was identified when it displayed signs of extreme aggression, entering two homes and biting three people. As a containment measure, the dog was taken to a veterinarian, where rabies was suspected. The dog died on February 10, and the case was laboratory-confirmed on February 11. Following this case, outbreak control activities were conducted by the local health center staff from February 12 to February 15. The surrounding area revealed multiple organic waste dumps and the remains of dead animals, coupled with reports of numerous stray and deceased dogs from neighbors. Fifteen days later, a second case was reported 890 meters away (Fig 1). This second dog bit two random people on the street and the owner reported their dog to the health center. The area where the second case was found was already being monitored and vaccinated due to the activities related to the first positive case. The health inspector observed signs similar to the initial positive case, prompting the owner to choose to euthanize the dog due to suspected rabies, which was confirmed later.

**Table 1.**
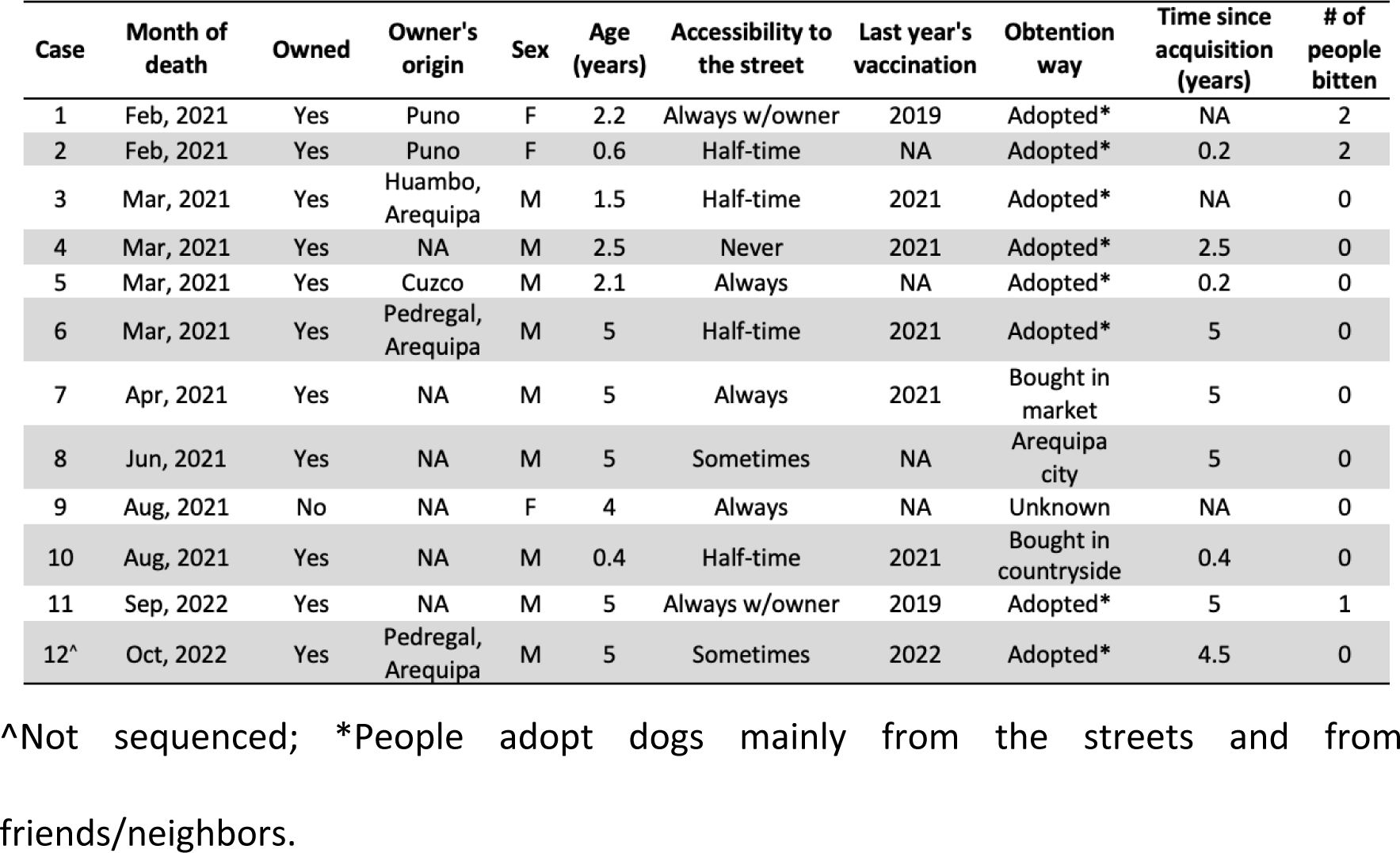
Epidemiological data collected during dog rabies control activities in El Pedregal.

Five days later, during outbreak control activities in the same area, the public health veterinarian from El Pedregal confirmed a third case; seven more cases were detected in the following weeks. Out of these 10 cases, eight were detected within a 1.95 km^2^ area, with the distance between cases ranging from 100 to 1,000 meters. In contrast, cases 8 and 9 were found approximately 3,300 m from the nearest case (Fig 1). Interestingly, six of the ten confirmed cases sought care in private veterinary clinics, where they received treatment for diseases other than rabies before notifying the public health center. Notably, the area where all the rabies cases were confirmed presented challenges for conducting outbreak control activities during regular working hours (i.e., 8 a.m. to 5 p.m.) due to the absence of owners engaged in agricultural work outside the city, returning home at night. To address this problem, health personnel conducted contact tracing and dog vaccinations during nocturnal hours. Two or three vaccination teams (each team comprising 1 vaccinator and 1 annotator) vaccinated up to 30 dogs daily, with the assistance of public health nurses for contact tracing and an ambulance for mobilization.

On September 7, 2022, 13 months following the cessation of reported cases, a new rabies incident surfaced in El Pedregal. A five-year-old dog was observed biting a cardboard box and subsequently bit a hen and a 10-year-old girl. Prompt intervention ensued as the family sought the assistance of a trusted veterinarian who aided in washing the girl’s wound. Following the veterinarian’s recommendation, the dog was taken to the public health center for evaluation. The bite was classified as mild, and rabies vaccination and antirabies serum were started. Concurrently, the public health zoonosis office at El Pedregal was notified, and at 9:30 am the dog was sent to a veterinary clinic for observation and died two hours later. A brain sample was extracted and dispatched to the referral laboratory in Arequipa City. On September 8, 2022, 33 dogs were vaccinated as part of the presumptive outbreak control measures. The case was not confirmed (by direct Fluorescent Antibody Test (DFAT), see ‘Routine Surveillance in Arequipa’) until September 13, with subsequent control activities prevented by logistical hurdles and owners’ absence during conventional working hours.

The final instance of rabies in El Pedregal was reported on October 17, 2022, when the owner of the affected dog reported it to the public health center. A private veterinarian, who had been administering the dog treatment for canine distemper since October 15, advised the owner to report the dog after observing neurological signs such as agitation, paralysis, and lethargy. Despite awareness of the mass dog vaccination campaign conducted in July 2022, the owner cited work commitments as hindering the pet’s vaccination, with the dog last receiving rabies vaccination in 2020. Suspecting rabies, the public health center veterinarian euthanized the dog on October 17, took a brain sample, and sent it to the laboratory on the same day, receiving DFAT confirmation of the case on October 18. The distance between these two cases from 2022 was 9,500 m Subsequent outbreak control measures were implemented on October 19, including vaccination of 24 dogs within the affected vicinity.

### Virus samples

A total of 34 virus samples from DFAT-confirmed rabid dogs in Peru (Arequipa, El Pedregal, Puno) were analyzed in this study. These samples, including 11 outbreak cases from El Pedregal, were obtained from different periods/sources and were sequenced in three distinct subsets. The following describes the samples in the order they were processed:

#### 1. Archived RNA from Puno (n=4, 2010-2012)

Before conducting sequencing in Peru, archived RNA from three dog-associated rabies cases (in livestock) sampled in 2011 and 2012 in Puno, a neighboring rabies-endemic region, were sequenced to obtain whole genome sequences (WGS) using an Illumina metagenomic approach at the Medical Research Council–University of Glasgow Centre for Virus Research (UoG-CVR), Glasgow, UK. These samples were collected from cattle with clinical signs of rabies as part of the routine surveillance activities of the National Service for Agrarian Health of Peru (SENASA). RNA extractions had previously been partially sequenced as part of a vampire bat rabies surveillance project (GenBank accession nos: KU938752 & KU938829) (25). The genomes produced here were utilized to design primers for dog variant RABV in Peru and in later phylogenetic analyses. An additional archived RNA sample from Puno in 2010 obtained from the same source was also sequenced at a later date, as part of batch 2 described below.

#### 2. Routine Surveillance in Arequipa (n=5, 2015-2019)

Five samples collected from routine rabies surveillance in Arequipa between 2015 and 2019 were included in the analysis. For diagnosis, whole brains were extracted and sent at room temperature to the Regional Reference Laboratory of Arequipa in a glycerin-saline solution transport medium. The presence of rabies virus antigen was evaluated using DFAT and confirmed by the mouse inoculation test and RT-PCR at the National Health Institute, following national regulations for rabies diagnosis (26). These samples were sequenced using an Illumina metagenomic approach at the UoG-CVR, UK.

#### 3. Outbreak Surveillance in El Pedregal (n=11, 2021-2022) and Ongoing Routine Surveillance in Arequipa (n=14, 2021-2023)

Samples from 11 rabies cases from the El Pedregal outbreak were obtained through local surveillance in this region (between February 2021 – September 2022, Fig 1). These samples underwent the same diagnostic procedures as described above and were sequenced using nanopore sequencing in the Zoonotic Disease Research Laboratory in Arequipa, Peru. In addition, 14 samples from Arequipa City, collected during the period 2021-2023, were sequenced using the same method.

### Multiplex primer scheme

Whole genomes from the three Puno RABV cases from 2011-12 (sample set 1) were used as references to design a multiplex primer scheme in Primal Scheme (27). Settings were applied to generate 400bp products with a 50bp overlap spanning the entirety of the genome.

### Nanopore sequencing

An established sample-to-sequence protocol was used to extract RNA from brain tissue and generate DNA libraries for nanopore sequencing (28). In brief, RNA was extracted and purified from homogenized brain tissue using a Zymo Quick RNA Miniprep kit with on-column DNase digestion (Zymo Research, USA). A 2-step RT-PCR was performed with RNA reverse transcribed using Lunascript RT Supermix (New England Biolabs, UK) and a multiplex PCR reaction using Q5 High Fidelity Hot-Start DNA Polymerase (New England Biolabs, UK) and the Peru RABV specific multiplex primer set. Library preparation was performed using a Nanopore ligation sequencing kit, SQK-LSK109, with native barcoding kit, EXP-NBD104/NBD114 (Oxford Nanopore Technologies, UK). Negative controls were included in each run to monitor cross-contamination. The final library was loaded onto an R9.4.1 flow cell and sequenced on a MinION device with live basecalling. Reads were processed using the bioinformatics pipeline described in Bautista, et al, 2023 (28) to produce consensus sequences. Any amplicon-specific contamination was masked in the final consensus sequence.

### Whole genome phylogenetic analysis

The 34 WGS generated in this study were aligned with an outgroup sequence, the RABV-GLUE reference sequence for the Cosmopolitan Africa 4 clade (GenBank accession: KF154998), using MAFFT v7.520 (29) with default settings. Phylogenetic tree reconstruction was performed using FastTree v2.1.11 (30) with a gtr+gamma substitution model and local support values obtained using FastTree’s default Shimodaira-Hasegawa test method. The tree was rooted by the outgroup, annotated, and visualized in R (31) v4.3.2 with the ggtree package (32). Maps were plotted in R using packages rnaturalearth and sf.

### Contextual phylogenetic analysis

RABV-GLUE (6), a bioinformatics resource for RABV sequence data, was utilized to obtain an alignment and associated metadata of all publicly available canine (excluding bat variant clades) RABV sequences from Latin America (n=1384). Details of these sequences are available in the GitHub repository. The 34 whole genome sequences (WGS) generated in this study were added to this alignment using the ‘add to existing alignment’ function in MAFFT v7.520 (29), ensuring the alignment length was preserved (33). A phylogenetic tree was produced as described above, using FastTree with annotation/visualization in R, rooted by the same outgroup described above.

### Data and scripts

Sequences of RABV are available at the NCBI GenBank repository (see accession numbers below and Table S1). Peru shapefiles were sourced from the Peruvian National Institute of Statistics and Informatics (https://ide.inei.gob.pe/#capas). All other data and scripts are available in the associated GitHub repository: https://github.com/RabiesLabPeru/Pedregal_genomics_outbreak_2021_2022.

## Results

We performed a phylogenetic characterization of a dog-mediated rabies outbreak in El Pedregal, analyzing 34 whole genome sequences (WGS) generated in this study—11 from El Pedregal and 23 from surrounding areas. Additionally, we included a contextual analysis with publicly available dog variant RABV sequences (>200bp) from the rest of Latin America. This study represents the first whole-genome analysis of dog-mediated RABV in Peru, and indeed in Latin America, with our sequences being the first dog variant rabies virus whole genomes from Peru to be published as available data in the GenBank repository (accession numbers PP965343-PP965374; existing records KU938752 & KU938829 updated to WGS). Comprehensive details, including sequencing platforms, library method, depths of coverage, and epidemiological data such as location, for all new sequences are provided in Table S1.

### Regional context of study sequences in Latin America

The current nomenclature for RABV phylogenetic classification is major clade, minor clade, and then lineage, representing increasingly finer genetic resolution (see (6) for detail). On a global scale, Peru RABV sequences, including those from Pedregal and Arequipa city, belong to the Cosmopolitan (Cosmo) major clade. A contextual analysis, including all available RABV sequences (of any length and from any genome region) from Latin America and excluding bat-variant or bat-derived (RAC-SK) sequences, was used to explore the relationship between RABVs from Peru and neighboring countries in detail. Within the Cosmo clade, Peruvian sequences cluster within a large section of the tree that lacks minor clade definition (annotated Cosmo:NA, Fig 2) but clearly stands apart from other known minor clades observed in neighboring countries, delineating at least one new minor clade that we have designated “Cosmopolitan America 5” (Cosmo:Am5, see annotation in Fig2A). Cosmo:Am5 also includes partial genome sequences (<1354bp) from Peru (35 sequences from the period 1985-2012) not from this study, and from Argentina (n=36), Bolivia (n=79) and Brazil (n=5) (Fig 2). The majority (78%) of these partial sequences are very short fragments (∼300bp), with only 26 providing full gene (nucleoprotein) level coverage. Note this excludes a portion of Cosmo:NA sequences from a related part of the tree, below the annotated Cosmo:Am5, that are genetically and geographically distinct (predominantly from island nations).

**Fig 2.**
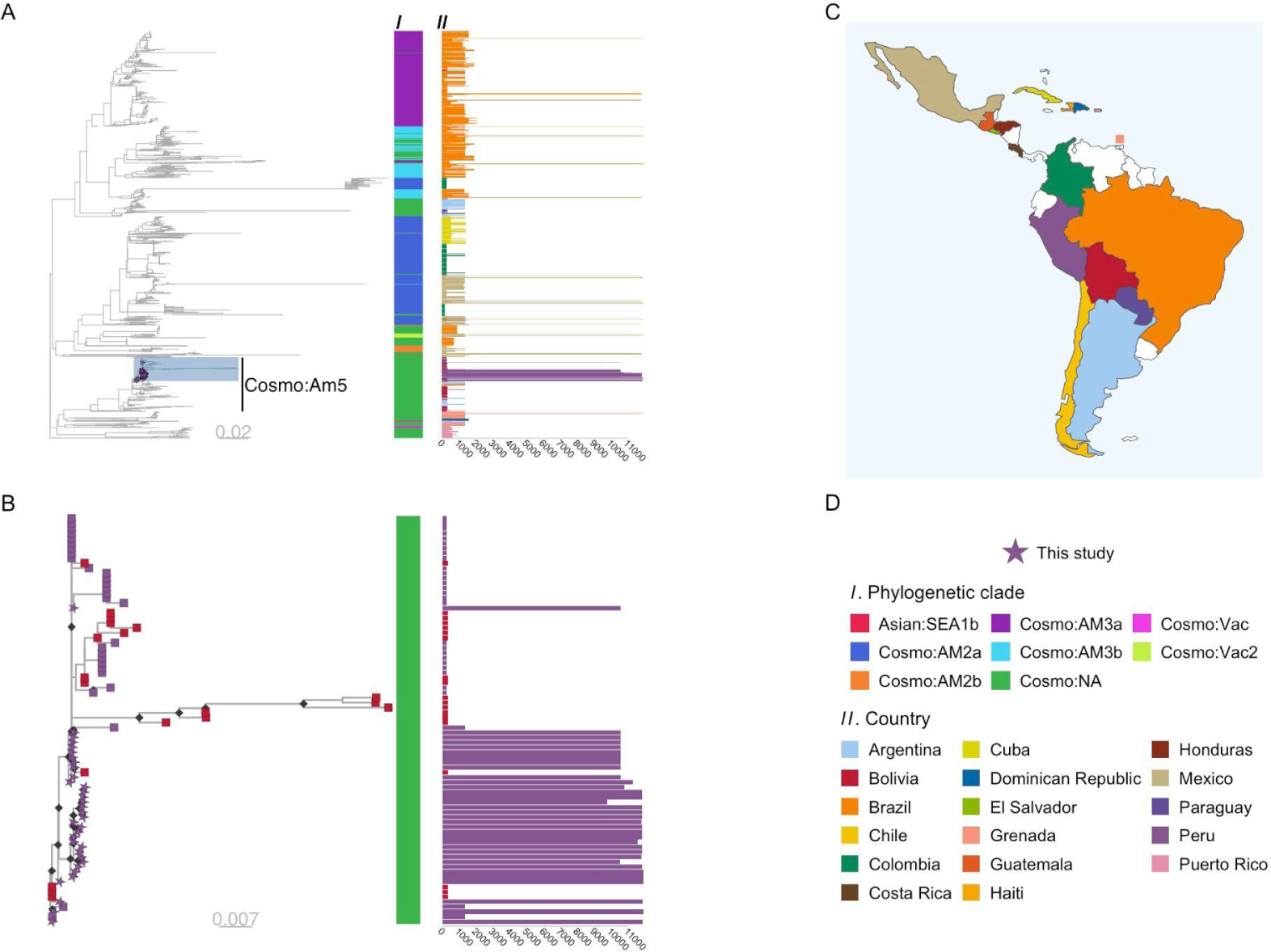
Phylogenetic trees of rabies virus (RABV) sequences from Latin America & the Caribbean (LAC) and newly sequenced genomes from Peru. Scaled in substitutions per site. (A) Phylogenetic tree of 1418 RABV sequences from LAC available in NCBI, which include sequences of any length and from any genome region, as well as 34 newly sequenced genomes from El Pedregal, Arequipa, and Puno in Peru. The tree is rooted by an outgroup sequence (GenBank accession: KF154998), not shown. The sequences from this study are highlighted and new minor clade Cosmo:Am5 is annotated. Color bar I indicates the phylogenetic clade of each sequence and the adjunct bar plot II shows sequence length (base pairs), colored by country of origin; (B) Subtree of the highlighted portion of tree A, showing all descendants of the most recent common ancestor of the Peruvian genomes sequenced in this study and related sequences. Tips are colored according to country of origin, with genomes from this study shown as stars. Color bar I and bar plot II follow the same scheme as in panel A; (C) Map of LAC with the countries of origin for the sequences indicated; (D) Colour schemes and annotation details.

Within the Cosmo:Am5 group there are further phylogenetic delineations between sequences that indicate the presence of multiple lineages, with possible geographic associations. The subtree of the most recent common ancestor (MRCA) of the Peruvian genomes from this study is expanded in Fig.2B and also contains descendants from Bolivia.

### El Pedregal phylogenetic outbreak analysis

The Peruvian WGS generated in this study were used to reconstruct a maximum-likelihood phylogeny. There was insufficient temporal signal in the data to enable a molecular clock-based analysis. The samples from Arequipa and Pedregal formed two distinct phylogenetic clusters (Fig 3), each predominantly containing sequences exclusively from their respective area. Ten of the eleven Pedregal sequences collected in 2021 formed one cluster sharing a common ancestor with one Arequipa sequence from 2019, which sits on an orphan branch ancestral to the Pedregal-only cases. The remaining Pedregal sample, representing the 2022 outbreak, clustered amongst all other samples from Arequipa (19 sequences spanning 2018 to 2023).

**Fig 3.**
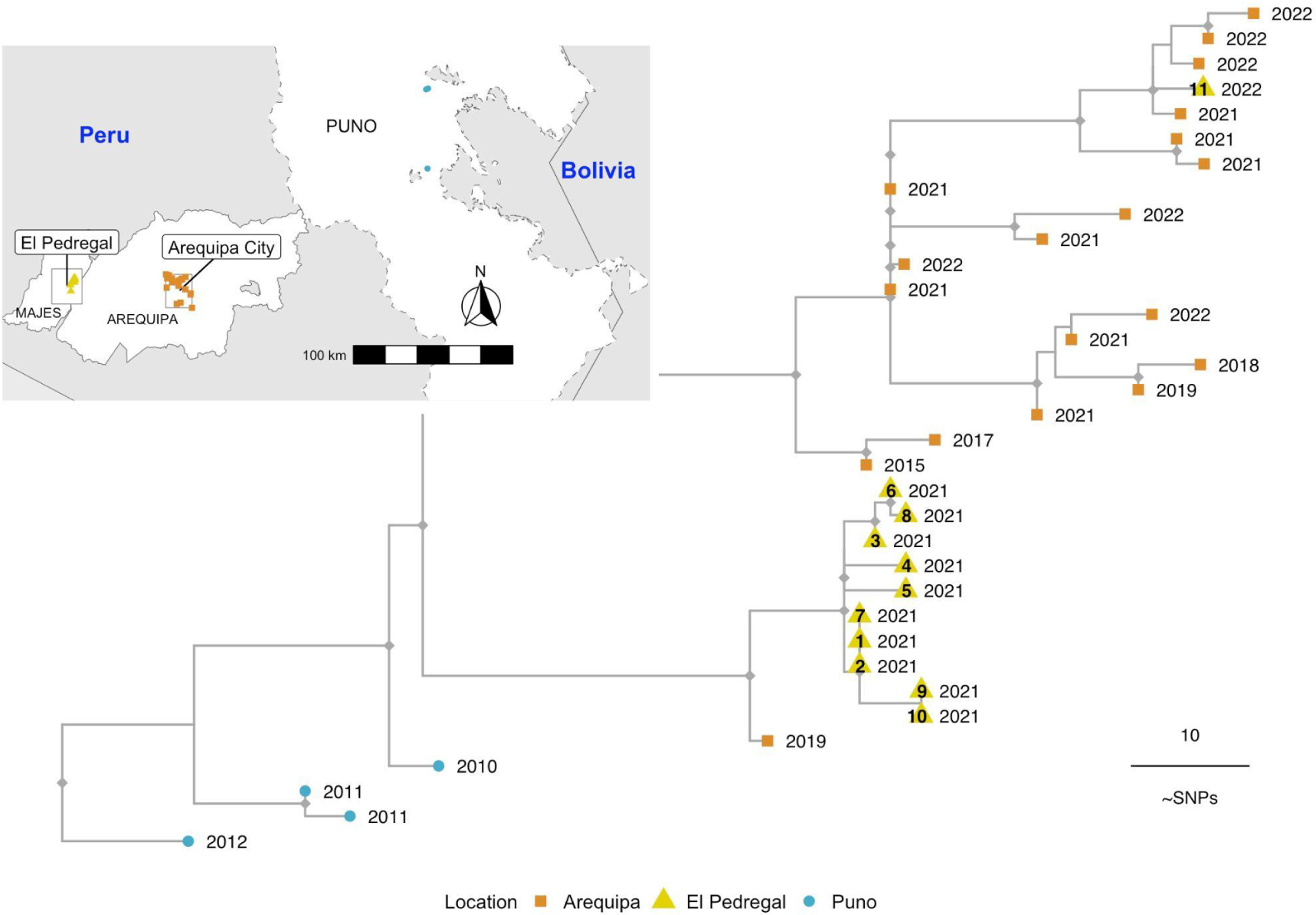
Phylogenetic tree of rabies virus (RABV) whole genome sequences from El Pedregal and surrounding areas. Tips are colored by geographic source, corresponding to the inset map, and labeled with the year of each RABV case. Sequences from El Pedregal (yellow triangles) are labeled with their case identifiers, reflecting the epidemiological timeline of cases and corresponding to Figures 1 and 4 as well as Table 1. The tree is rooted with an outgroup sequence (GenBank accession: KF154998, not shown) and the scale represents the number of mutations (SNPs: single nucleotide polymorphisms).

These results suggest that the El Pedregal 2021 outbreak resulted from a single introduction and brief establishment of local RABV transmission rather than repeated introductions. Given the limited data available, the source of this introduction remains unclear. However, its distinction from Arequipa City cases (except for one isolated case in Arequipa; Fig 3) suggests that it did not come directly from Arequipa but rather that both areas share a common source of introduction, which, considering the ancestral evolutionary history (Fig 3) alongside the epidemiological and demographic context, is likely to be Puno region. In contrast, the El Pedregal 2022 outbreak was caused by a new introduction clearly originating from Arequipa City.

### Epidemiological analysis

We used the epidemiological data collected during outbreak control activities to reconstruct the timeline of cases in El Pedregal (Fig 4). We inferred a likely transmission chain between case number one with cases two and three (owners were relatives who visited each other with their dogs and lived close to each other); and case number three with cases four and six (owners were acquaintances who visited each other with their dogs). In 2021, seven out of ten cases occurred within the first three months after the index case. It is important to mention that the year before this outbreak the mass vaccination campaign in El Pedregal was canceled due to the pandemic. Thus, all rabid dogs were not vaccinated in the previous year. Interestingly, five of them were vaccinated during the outbreak control activities (Fig 1) but still developed rabies. Among these dogs, the time from vaccination to showing rabies signs was 14, 13, 56, 148, and 1 days. Additionally, the reconstructed timeline shows an epidemiological silence of one year from the last rabid dog reported in 2021 until the next case in 2022 which was the result of a second introduction to El Pedregal based on the genetic analysis.

**Fig 4.**
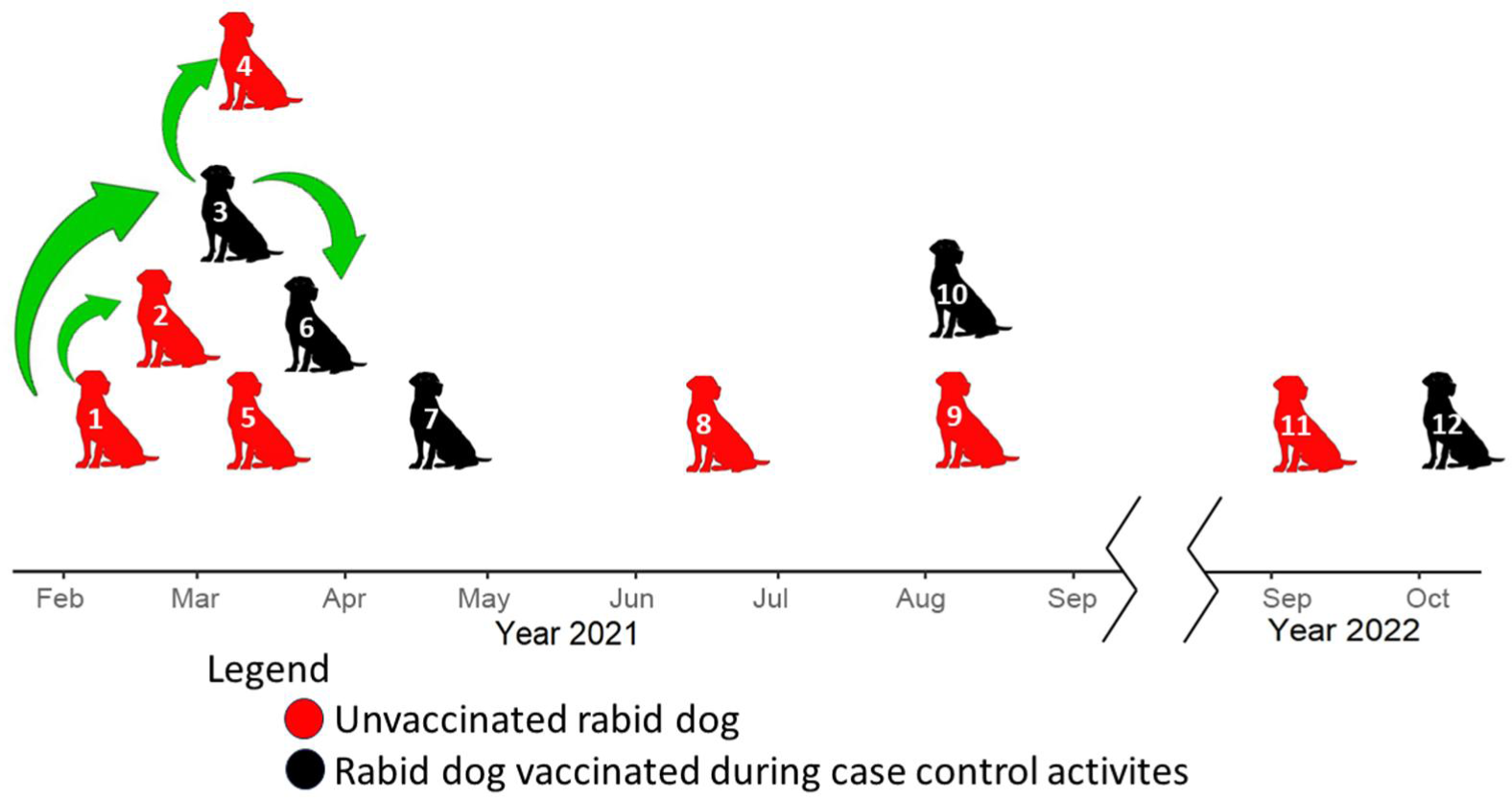
Timeline of dog rabies outbreaks in El Pedregal. The figure highlights the relationship between the cases according to epidemiological data and the reported vaccination status of the rabid dogs.

The epidemiological data (Table 1), reveals that most rabid dogs were owned (11/12) and had regular access to the outdoors (at least some hours a week outdoors without restriction) (9/12). Three out of 6 owners were from Puno, and only one was from El Pedregal; we could not obtain this information from the other owners. Dogs’ ages ranged from five months to five years, and three dogs were obtained or adopted from the streets during the outbreak. Additionally, people who were bitten sought treatment for their wounds at the health center between 4 to 7 days after the bite.

## Discussion

In this study, we investigate a dog rabies outbreak in El Pedregal, Arequipa using detailed epidemiological and genomic data.

Our findings reveal that the canine RABV circulating in southern Peru belongs to a novel minor clade within the Cosmopolitan major clade, which we have designated “Cosmopolitan Am5,” in accordance with existing RABV phylogenetic nomenclature (34,35). This minor clade is distinct from other canine RABV minor clades previously described in LAC, including those in neighboring countries. However, our analysis indicates that this clade has been present on the continent since at least 1985, with the first sequence detection in a domestic dog from Lima, Peru (NCBI accession: KF831564).

Notably, most of the sequences from this new minor clade, and, LAC sequences in general, are only partial gene or gene level coverage sequences except for the new genomes produced in this study. This limitation makes comparative phylogenetic analyses challenging and restricts the ability to fully utilize genomic data to analyze canine RABV diversity, its circulation and spread, and to pinpoint external sources of introduction. This lack of sequence data and lack of investment in WGS is likely due to the significant progress in dog rabies elimination in LAC over the last 40 years (36), which coincided with the rise of genome sequencing accessibility and its use as a key surveillance tool (11,37). Despite these challenges, our analyses demonstrate the value and potential of genomics-informed surveillance to inform dog rabies outbreak response in LAC. Even with a small WGS dataset from our study area, we were able to perform a contextual analysis to identify potential connections with other LAC countries and a fine-scale local analysis that ruled out Arequipa (the most obvious hypothesis) as the source of the initial outbreak in El Pedregal. Furthermore, this sets a foundation for future genomic surveillance work to help understand and eliminate the remaining pockets of endemic dog rabies in LAC.

Our epidemiological data from the 2021 dog RABV outbreak in El Pedregal suggest that two secondary cases can be linked to the index case; one of these secondary cases also produced two new cases of its own. Our WGS phylogenetic analysis supports a local transmission dynamic, showing that all cases sequenced during this period (n=10) clustered together, sharing a common ancestor, and were genetically distinct from the dog RABV circulating in Arequipa City. This supports the hypothesis that a single introduction, followed by local transmission in El Pedregal, was responsible for the 2021 outbreak. The Andean desert around El Pedregal does not harbor any species likely to act as reservoirs of rabies in the area, making human-mediated translocation of infected dogs the most probable mechanism of rabies introductions to El Pedregal.

While our analysis was unable to pinpoint the exact source of introduction for the 2021 outbreak, it was clear from the available data that it did not come directly from Arequipa City, despite a large ongoing dog rabies outbreak in the city for the last eight years and its proximity and continued population interchange with El Pedregal (17,38). However, a second outbreak in 2022 in El Pedregal did appear to result from an introduction from Arequipa City, rather than the persistence of the first outbreak. Hence, even with limited sequence data El Pedregal demonstrates evidence of at least two introductions within two years, underscoring that introduction and re-emergence is a persistent threat in the region, despite its geographic isolation surrounded by desert. Arequipa City and the Puno region are both potential sources of introduction as neighboring and endemic areas that report active cases of dog rabies (12,39). Furthermore, the emergence of RABV in Arequipa has previously been linked back to an introduction from Puno (40).

Our analysis is limited by the small number of rabies cases and sequences available from this outbreak. There were also relatively few historical RABV sequences from Latin America (11), and they were mainly restricted to very short sequences (e.g. partial gene ∼200-300bp), limiting the degree to which we could infer outbreak origins. For future studies, we suggest obtaining genomes from a larger number of samples from Arequipa City and the Puno region over the same period to provide more conclusive evidence of the source of RABV emergence in El Pedregal and facilitate more comprehensive phylodynamic analyses that could elucidate transmission drivers and diffusion pathways.

Across diverse global regions where dog RABV has been studied, incursions of dog RABV appear to be common, with genomic surveillance revealing higher rates than expected (9,21). Human-mediated movement of dogs has emerged as a significant driver (41,42) of these occurrences. While many introductions fail to establish due to stochastic factors in rabies transmission (43), areas with increased human movement are likely to face heightened risks. This is evident in El Pedregal, which has a history of migratory settlement and where a portion of the population commutes daily, weekly, or seasonally from nearby cities to Pedregal for work (17). Moreover, poor housing conditions in the peri-urban areas of El Pedregal prevent dog owners from keeping dogs inside homes, making these dogs less accessible and harder to vaccinate, and increasing the vulnerability of El Pedregal to new introductions, similar to the ongoing rabies outbreak in Arequipa City (15,17). In Arequipa City, such conditions were identified as the main factors that allowed the introduction and persistence of the RABV, together with landscape features, such as dry water channels that allowed unrestricted and fast movement of dogs across large parts of the city, facilitating the rapid spread of the RABV (15,44).

Mass dog vaccination will be crucial to preventing introductions and their onward spread. The year before the 2021 outbreak in El Pedregal, mass dog vaccinations were canceled due to the COVID-19 pandemic, providing a window of opportunity for the virus to take hold. Although the outbreak prompted a rapid local response, including dog vaccinations, five of the vaccinated dogs still contracted and died from rabies, even after months of being vaccinated. For most cases, it remains unclear whether the vaccine was administered too late or if it was ineffective. Assessing the efficacy of these ring vaccination activities is crucial. Currently, in Arequipa, when owners of exposed dogs do not permit euthanasia, these dogs are vaccinated in an attempt to prevent new cases. Furthermore, the last reported rabies case in El Pedregal (October 17, 2022) did not receive vaccination during the 2022 outbreak response but had been vaccinated in 2020, according to its owner. While annual boosters are recommended, these observations raise questions about the vaccine efficacy or the administration process (e.g. poorly trained vaccinators might inoculate in the wrong area/tissue).

Genomic information has the potential to shed light on complex epidemiological scenarios. This study provides a snapshot of RABV introduction in a rapidly urbanizing rural area, a common scenario in Latin American cities. Our results indicate multiple introductions into El Pedregal in 2021 and 2022, most likely mediated by human translocation of dogs. This study highlights the potential of genomics analysis to understand rabies outbreaks, enhance surveillance systems, and inform rabies control efforts.

## Acknowledgments

We gratefully acknowledge the Gerencia Regional de Salud de Arequipa and Red de Salud Arequipa Caylloma who conducted the focus control activities and shared their expertise with our team. We are grateful to the National Service of Agrarian Health (SENASA) of Peru for providing data on rabies incidence in livestock and for allowing access to rabies positive samples for whole genome sequencing and thanks to Daniel Streicker and Alice Broos for providing extracted RNA from archived samples. We thank Antuannete Vela for her technical support with case reconstruction information and Guillermo Porras and Paul Tevez for supporting the focus control activities.

## Supplementary files

**S1 Table.** Sequencing and epidemiology details of newly sequenced rabies virus sequences used in this study.

| NCBI accession | Isolate ID | Case ID | NGS platform | Library type | Virus sample set | NGS run | Mapped reads | Mean (stdev) coverage | Sequence length* | Sample Collection Date | Outbreak place | Region | Province | District |
| --- | --- | --- | --- | --- | --- | --- | --- | --- | --- | --- | --- | --- | --- | --- |
| PP965373 | 1101786 | - | Illumina | Metagenomic | 1 | illumina-1 | 173909 | 2987 (741) | 11869 | 26-Mar-11 | Puno | Puno | Azangaro | Caminaca |
| KU938752* | 1101787 | - | Illumina | Metagenomic | 1 | illumina-1 | 731825 | 6536 (1471) | 11874 | 3-Mar-11 | Puno | Puno | Azangaro | Caminaca |
| KU938829* | 1203037 | - | Illumina | Metagenomic | 1 | illumina-1 | 12966 | 237 (169) | 11842 | 8-May-12 | Puno | Puno | Puno | Atuncolla |
| PP965367 | 107_2018 | - | Illumina | Metagenomic | 2 | illumina-2 | 1955 | 47 (52) | 10500 | 25-Mar-18 | Arequipa | Arequipa | Arequipa | Cerro Colorado |
| PP965364 | 107_2021 | 6 | Nanopore | Amplicon | 3 | nano-1 | 360339 | 13521 (7447) | 10563 | 20-Mar-21 | El Pedregal | Arequipa | Caylloma | Majes |
| PP965360 | 121_2019 | - | Illumina | Metagenomic | 2 | illumina-2 | 1236 | 30 (12) | 11791 | 22-Apr-19 | Arequipa | Arequipa | Arequipa | Cerro Colorado |
| PP965366 | 126_2021 | - | Nanopore | Amplicon | 3 | nano-2 | 41076 | 1826 (726) | 11816 | 31-Mar-21 | Arequipa | Arequipa | Arequipa | Cerro Colorado |
| PP965361 | 145_2022 | - | Nanopore | Amplicon | 3 | nano-2 | 69588 | 2757 (1405) | 11827 | 9-Jun-22 | Arequipa | Arequipa | Arequipa | Cerro Colorado |
| PP965351 | 147_2021 | 7 | Nanopore | Amplicon | 3 | nano-1 | 372481 | 14426 (6599) | 10565 | 19-Apr-21 | El Pedregal | Arequipa | Caylloma | Majes |
| PP965365 | 147_2022 | - | Nanopore | Amplicon | 3 | nano-2 | 35162 | 1666 (1144) | 10752 | 10-Jun-22 | Arequipa | Arequipa | Arequipa | Yura |
| PP965369 | 158_2022 | - | Nanopore | Amplicon | 3 | nano-2 | 26045 | 1500 (545) | 11825 | 21-Jun-22 | Arequipa | Arequipa | Arequipa | Jacobo Hunter |
| PP965371 | 172_2022 | - | Nanopore | Amplicon | 3 | nano-2 | 83048 | 3167 (1618) | 11812 | 8-Jul-22 | Arequipa | Arequipa | Arequipa | Cayma |
| PP965368 | 173_2022 | - | Nanopore | Amplicon | 3 | nano-2 | 60646 | 2606 (1257) | 11819 | 12-Jul-22 | Arequipa | Arequipa | Arequipa | Jacobo Hunter |
| PP965362 | 182_2019 | - | Illumina | Metagenomic | 2 | illumina-2 | 1044 | 25 (16) | 11273 | 16-Jun-19 | Arequipa | Arequipa | Arequipa | Alto Selva Alegre |
| PP965344 | 202_2021 | 8 | Nanopore | Amplicon | 3 | nano-1 | 112087 | 7031 (1900) | 10563 | 14-Jun-21 | El Pedregal | Arequipa | Caylloma | Majes |
| PP965347 | 219_2022 | - | Nanopore | Amplicon | 3 | nano-2 | 36608 | 2472 (878) | 11822 | 31-Aug-22 | Arequipa | Arequipa | Arequipa | Characato |
| PP965374 | 231_2021 | - | Nanopore | Amplicon | 3 | nano-2 | 28192 | 1405 (515) | 11801 | 15-Jul-21 | Arequipa | Arequipa | Arequipa | Cerro Colorado |
| PP965345 | 236_2022 | 12 | Nanopore | Amplicon | 3 | nano-1 | 758917 | 25058 (17387) | 9755 | 7-Sep-22 | El Pedregal | Arequipa | Caylloma | Majes |
| PP965363 | 250_2021 | 9 | Nanopore | Amplicon | 3 | nano-1 | 350173 | 14068 (5935) | 10565 | 10-Aug-21 | El Pedregal | Arequipa | Caylloma | Majes |
| PP965354 | 251_2021 | 10 | Nanopore | Amplicon | 3 | nano-1 | 115094 | 7740 (2039) | 10565 | 10-Aug-21 | El Pedregal | Arequipa | Caylloma | Majes |
| PP965359 | 30_2021 | - | Nanopore | Amplicon | 3 | nano-2 | 50742 | 2183 (936) | 11821 | 5-Feb-21 | Arequipa | Arequipa | Arequipa | Yura |
| PP965357 | 31_2021 | - | Nanopore | Amplicon | 3 | nano-2 | 71634 | 2593 (1262) | 11820 | 8-Feb-21 | Arequipa | Arequipa | Arequipa | Cerro Colorado |
| PP965349 | 34_2021 | 1 | Nanopore | Amplicon | 3 | nano-1 | 148926 | 7481 (2403) | 10565 | 10-Feb-21 | El Pedregal | Arequipa | Caylloma | Majes |
| PP965355 | 3553_2010 | - | Illumina | Metagenomic | 1 | illumina-2 | 3848 | 95 (52) | 11904 | 2010 | Puno | Puno | Azangaro |  |
| PP965352 | 375_2017 | - | Illumina | Metagenomic | 2 | illumina-2 | 3545 | 88 (82) | 11893 | 23-Nov-17 | Arequipa | Arequipa | Arequipa | Mariano Melgar |
| PP965353 | 40_2021 | - | Nanopore | Amplicon | 3 | nano-2 | 9237 | 393 (169) | 11826 | 15-Feb-21 | Arequipa | Arequipa | Arequipa | Cerro Colorado |
| PP965343 | 52_2021 | 2 | Nanopore | Amplicon | 3 | nano-1 | 47951 | 3833 (1027) | 10565 | 24-Feb-21 | El Pedregal | Arequipa | Caylloma | Majes |
| PP965358 | 56_2021 | - | Nanopore | Amplicon | 3 | nano-2 | 70889 | 2336 (1445) | 11580 | 27-Feb-21 | Arequipa | Arequipa | Arequipa | Yura |
| PP965348 | 560_2015 | - | Illumina | Metagenomic | 2 | illumina-2 | 2106 | 52 (30) | 11884 | 7-Sep-15 | Arequipa | Arequipa | Arequipa | Mariano Melgar |
| PP965346 | 61_2021 | - | Nanopore | Amplicon | 3 | nano-2 | 70413 | 2947 (1222) | 11826 | 1-Mar-21 | Arequipa | Arequipa | Arequipa | Cerro Colorado |
| PP965356 | 68_2021 | 3 | Nanopore | Amplicon | 3 | nano-1 | 245893 | 10765 (4169) | 10563 | 3-Mar-21 | El Pedregal | Arequipa | Caylloma | Majes |
| PP965350 | 71_2021 | 4 | Nanopore | Amplicon | 3 | nano-1 | 328914 | 13188 (5130) | 10564 | 6-Mar-21 | El Pedregal | Arequipa | Caylloma | Majes |
| PP965370 | 73_2021 | 5 | Nanopore | Amplicon | 3 | nano-1 | 42452 | 3467 (812) | 10564 | 6-Mar-21 | El Pedregal | Arequipa | Caylloma | Majes |
| PP965372 | 90_2021 | - | Nanopore | Amplicon | 3 | nano-2 | 45757 | 1838 (825) | 11826 | 13-Mar-21 | Arequipa | Arequipa | Arequipa | Yura |

\*Genbank records updated

## References

1. Fahrion AS, Mikhailov A, Giacinti J, Harries J. Human rabies transmitted by dogs: current status of global data, 2015. Weekly Epidemiological Record. 2016;(2).

2. Hampson K, Coudeville L, Lembo T, Sambo M, Kieffer A, Attlan M, et al. Estimating the Global Burden of Endemic Canine Rabies. Carvalho MS, editor. PLoS Negl Trop Dis. 2015 Apr 16;9(4):e0003709.

3. World Health Organization, Food and Agriculture Organization of the United Nations, World Organisation for Animal Health, Global Alliance for Rabies Control. United Against Rabies Collaboration. First annual progress report: Global Strategic Plan to End Human Deaths from Dog-mediated Rabies by 2030. Geneva; 2019.

4. Brunker K, Lemey P, Marston DA, Fooks AR, Lugelo A, Ngeleja C, et al. Landscape attributes governing local transmission of an endemic zoonosis: Rabies virus in domestic dogs. Mol Ecol. 2018 Feb;27(3):773–88.

5. Marston DA, Horton DL, Nunez J, Ellis RJ, Orton RJ, Johnson N, et al. Genetic analysis of a rabies virus host shift event reveals within-host viral dynamics in a new host. Virus Evolution [Internet]. 2017 Jul 1 [cited 2024 Jun 18];3(2). Available from: https://academic.oup.com/ve/article/doi/10.1093/ve/vex038/4737086

6. Campbell K, Gifford RJ, Singer J, Hill V, O’Toole A, Rambaut A, et al. Making genomic surveillance deliver: A lineage classification and nomenclature system to inform rabies elimination. Lauring AS, editor. PLoS Pathog. 2022 May 2;18(5):e1010023.

7. Dellicour S, Troupin C, Jahanbakhsh F, Salama A, Massoudi S, Moghaddam MK, et al. Using phylogeographic approaches to analyse the dispersal history, velocity and direction of viral lineages — Application to rabies virus spread in Iran. Molecular Ecology. 2019 Sep;28(18):4335–50.

8. Gibson AD, Yale G, Corfmat J, Appupillai M, Gigante CM, Lopes M, et al. Elimination of human rabies in Goa, India through an integrated One Health approach. Nat Commun. 2022 May 19;13(1):2788.

9. Lushasi K, Brunker K, Rajeev M, Ferguson EA, Jaswant G, Baker LL, et al. Integrating contact tracing and whole-genome sequencing to track the elimination of dog-mediated rabies: An observational and genomic study. eLife. 2023 May 25;12:e85262.

10. Brunker K, Nadin-Davis S, Biek R. Genomic sequencing, evolution and molecular epidemiology of rabies virus. Rev Sci Tech Off Int Epizoot. 2018 Aug 1;37(2):401– 8.

11. Jaswant G, Bautista CT, Ogoti B, Changalucha J, Oyugi JO, Campbell K, et al. Viral sequencing to inform the global elimination of dog-mediated rabies - a systematic review. One Health Implement Res. 2024 May 31;4(2):15–37.

12. Organización Panamericana de la Salud. Eliminacion de la rabia humana transmitida por peros en America Latin: Análisis de la situación. Washington, D.C.: Organizacion Panamericana de la Salud; 2005.

13. Vigilato MAN, Clavijo A, Knobl T, Silva HMT, Cosivi O, Schneider MC, et al. Progress towards eliminating canine rabies: policies and perspectives from Latin America and the Caribbean. Phil Trans R Soc B. 2013 Aug 5;368(1623):20120143.

14. Dirección General de Epidemiología. Alerta Epidemiológica: Alerta ante la identificación de casos de rabia canina en Arequipa y riesgo de rabia urbana humana. Lima, Perú: Ministerio de Salud del Perú (MINSA); 2015 Feb. (Alerta Epidemiológica). Report No.: Report No.: AE-DEVE N. 003-2015.

15. Castillo-Neyra R, Brown J, Borrini K, Arevalo C, Levy MZ, Buttenheim A, et al. Barriers to dog rabies vaccination during an urban rabies outbreak: Qualitative findings from Arequipa, Peru. Recuenco S, editor. PLoS Negl Trop Dis. 2017 Mar 17;11(3):e0005460.

16. Raynor B, Díaz EW, Shinnick J, Zegarra E, Monroy Y, Mena C, et al. The impact of the COVID-19 pandemic on rabies reemergence in Latin America: The case of Arequipa, Peru. Blanton J, editor. PLoS Negl Trop Dis. 2021 May 21;15(5):e0009414.

17. Gonçalves R, Hacker KP, Condori C, Xie S, Borrini-Mayori K, Mollesaca Riveros L, et al. Irrigation, migration and infestation: a case study of Chagas Disease Vectors and bed bugs in El Pedregal, Peru. Mem Inst Oswaldo Cruz. 2024;119:e240002.

18. Erwin A, Ma Z, Popovici R, Salas O’Brien EP, Zanotti L, Silva CA, et al. Linking migration to community resilience in the receiving basin of a large-scale water transfer project. Land Use Policy. 2022 Mar;114:105900.

19. Municipalidad Distrital de Majes. Campaña gratuita de vacunación antirrábica canina (Comunicado N° 061-2023/UIIYRP/MDM) [Internet]. 2023 [cited 2023 Jul 4]. Available from: https://www.gob.pe/institucion/munimajes/noticias/788762-campana-gratuita-de-vacunacion-antirrabica-canina

20. Brunker K, Jaswant G, Thumbi SM, Lushasi K, Lugelo A, Czupryna AM, et al. Rapid in-country sequencing of whole virus genomes to inform rabies elimination programmes [version 2; peer review: 3 approved]. Wellcome Open Research. 2020;

21. Bourhy H, Nakouné E, Hall M, Nouvellet P, Lepelletier A, Talbi C, et al. Revealing the Micro-scale Signature of Endemic Zoonotic Disease Transmission in an African Urban Setting. Parrish C, editor. PLoS Pathog. 2016 Apr 8;12(4):e1005525.

22. Zuniga L. Transformation of the hyperarid desert soils in a Arequipa Peru during four decades of irrigation agriculture. [West Lafayette, Indiana]: Purdue University Graduate School; 2020.

23. Zapana Churata LE. Respuestas a la crisis hídrica en zonas agrícolas y urbanas: Caso de estudio “Proyecto de Irrigación Majes Siguas I” Arequipa – Perú. Agua Territorio. 2018 Nov 13;(12):145–56.

24. Instituto Nacional de Estadística e Informática. Peru: Estimaciones y proyeccciones de población por departamento, Provincia y Distrito 2018-2020 [Internet]. Lima, Perú; 2020 Enero [cited 2023 Jul 11]. (Boletin especial). Report No.: N0. 26. Available from: https://www.inei.gob.pe/media/MenuRecursivo/publicaciones_digitales/Est/Lib1715 /libro.pdf

25. Streicker DG, Winternitz JC, Satterfield DA, Condori-Condori RE, Broos A, Tello C, et al. Host–pathogen evolutionary signatures reveal dynamics and future invasions of vampire bat rabies. Proc Natl Acad Sci USA. 2016 Sep 27;113(39):10926–31.

26. Instituto Nacional de Salud. Manual de procedimientos para el diagnóstico de la rabia. Lima: Ministerio de Salud; 2002. 46 p.

27. Quick J, Loman NJ, Duraffour S, Simpson JT, Severi E, Cowley L, et al. Real-time, portable genome sequencing for Ebola surveillance. Nature. 2016 Feb;530(7589):228–32.

28. Bautista C, Jaswant G, French H, Campbell K, Durrant R, Gifford R, et al. Whole Genome Sequencing for Rapid Characterization of Rabies Virus Using Nanopore Technology. 2023; Available from: https://review.jove.com/t/65414/whole-genome-sequencing-for-rapid-characterization-rabies-virus-using?status=a67420k

29. Katoh K, Standley DM. MAFFT Multiple Sequence Alignment Software Version 7: Improvements in Performance and Usability. Molecular Biology and Evolution. 2013 Apr 1;30(4):772–80.

30. Price MN, Dehal PS, Arkin AP. FastTree 2 – Approximately Maximum-Likelihood Trees for Large Alignments. Poon AFY, editor. PLoS ONE. 2010 Mar 10;5(3):e9490.

31. R Core Team. R: A language and environment for statistical computing [Internet]. Vienna, Austria: R Foundation for Statistical Computing; 2021. Available from: URL https://www.R-project.org/

32. Xu S, Li L, Luo X, Chen M, Tang W, Zhan L, et al. *Ggtree* : A serialized data object for visualization of a phylogenetic tree and annotation data. iMeta. 2022 Dec;1(4):e56.

33. Katoh K, Frith MC. Adding unaligned sequences into an existing alignment using MAFFT and LAST. Bioinformatics. 2012 Dec 1;28(23):3144–6.

34. Troupin C, Dacheux L, Tanguy M, Sabeta C, Blanc H, Bouchier C, et al. Large-Scale Phylogenomic Analysis Reveals the Complex Evolutionary History of Rabies Virus in Multiple Carnivore Hosts. PLoS Pathog. 2016 Dec;12(12):e1006041.

35. Kuzmin IV, Shi M, Orciari LA, Yager PA, Velasco-Villa A, Kuzmina NA, et al. Molecular inferences suggest multiple host shifts of rabies viruses from bats to mesocarnivores in Arizona during 2001-2009. PLoS Pathog. 2012;8(6):e1002786.

36. Vigilato MAN, Belotto AJ, Tamayo Silva H, Rocha F, Molina-Flores B, Pompei JCA, et al. Towards the Elimination of Canine Rabies in the Americas: Governance of a Regional Program. In: Rupprecht CE, editor. History of Rabies in the Americas: From the Pre-Columbian to the Present, Volume I: Insights to Specific Cross-Cutting Aspects of the Disease in the Americas [Internet]. Cham: Springer International Publishing; 2023. p. 293–305. Available from: 10.1007/978-3-031-25052-1_13

37 . Gardy JL, Loman NJ. Towards a genomics-informed, real-time, global pathogen surveillance system. Nature Reviews Genetics. 2018 Jan 1;19(1):9–20.

38. Castillo-Neyra R, Toledo AM, Arevalo-Nieto C, MacDonald H, De La Puente-León M, Naquira-Velarde C, et al. Socio-spatial heterogeneity in participation in mass dog rabies vaccination campaigns, Arequipa, Peru. Blanton J, editor. PLoS Negl Trop Dis. 2019 Aug 1;13(8):e0007600.

39. Centro Nacional de Epidemiología, Prevención y Control de Enfermedades. Boletín Epidemiológico Semana Epidemiológica 51 (del 18 al 24 de diciembre 2022). Lima, Perú: Ministerio de Salud del Perú (MINSA); 2022. (Boletín Epidemiológico). Report No.: Volumen 31-SE 51-2022.

40. Mantari Torpoco CR, Berrocal Huallpa AM, Espinoza-Culupú AO, López-Ingunza RL. Caracterización molecular de la nucleoproteína del virus de la rabia en canes procedentes de Arequipa, Perú. Rev Peru Med Exp Salud Publica. 2019 Mar 21;36(1):46.

41. Townsend SE, Sumantra IP, Pudjiatmoko, Bagus GN, Brum E, Cleaveland S, et al. Designing programs for eliminating canine rabies from islands: Bali, Indonesia as a case study. PLoS Negl Trop Dis. 2013;7(8):e2372.

42. Brunker K, Marston DA, Horton DL, Cleaveland S, Fooks AR, Kazwala R, et al. Elucidating the phylodynamics of endemic rabies virus in eastern Africa using whole-genome sequencing. Virus Evol. 2015;1(1):vev011.

43. Hampson K, Dushoff J, Cleaveland S, Haydon DT, Kaare M, Packer C, et al. Transmission dynamics and prospects for the elimination of canine rabies. PLoS Biol. 2009 Mar 10;7(3):e53.

44. Castillo-Neyra R, Zegarra E, Monroy Y, Bernedo R, Cornejo-Rosello I, Paz-Soldan V, et al. Spatial Association of Canine Rabies Outbreak and Ecological Urban Corridors, Arequipa, Peru. TropicalMed. 2017 Aug 13;2(3):38.

